# Multidimensionality in the thermal niches of dung beetles could limit species’ responses to temperature changes

**DOI:** 10.1101/2020.11.15.383612

**Authors:** Joaquín Calatayud, Joaquín Hortal, Jorge Ari Noriega, Ángel Arcones, Verónica R. Espinoza, Noemí Guil, Jorge M. Lobo

**Affiliations:** Departamento de Biología y Geología, Física y Química inorgánica. Universidad Rey Juan Carlos, C/ Tulipán s/n, Móstoles, Madrid, 28933 Spain; Department of Biogeography and Global Change, Museo Nacional de Ciencias Naturales (MNCN□CSIC), C/José Gutiérrez Abascal, 2, Madrid, 28006 Spain; Laboratorio de Zoología y Ecología Acuática – LAZOEA, Universidad de los Andes, Bogotá, Colombia; Facultad de Medicina Veterinaria y Zootecnia. Universidad Central del Ecuador, Ecuador

**Keywords:** biological scale, daily activity, geographic distribution, niche dimensionality, phenology, physiological trade-offs

## Abstract

Understanding the consequences of climate change requires understanding how temperature controls species’ responses across key biological aspects, as well as the coordination of thermal responses across these aspects. We study the role of temperature in determining the species’ diel, seasonal, and geographical occurrence, using dung beetles as a model system. We found that temperature has relatively low −but not negligible− effects in the three spatiotemporal scales, once accounting for alternative factors. More importantly, the estimated thermal responses were largely incongruent across scales. This shows that species have multidimensional thermal niches, entailing that adjustments to fulfil temperature requirements for one biological aspect, such as seasonal ontogenetic cycles, may result in detrimental effects on other aspects, like diel activity. These trade-offs can expose individuals to inadequate temperatures, reducing populations’ performance. Paradoxically, the relatively weak effects of temperature we found may have serious consequences for species’ responses to warming if temperature regulates essential aspects of species’ biology in divergent ways.

## Introduction

Temperature is fundamental for the efficient capture and management of the energy that maintains living organisms (Brown et al. 2004). Temperature variations affect the abundance and distribution of species (Angilletta 2009), the variability of ecological systems (Wang et al. 2009), and even the history of life and biodiversity on Earth itself (Schwartzman 1999, Mayhew et al. 2008). Indeed, temperature plays a critical role in controlling key aspects such as species’ spatiotemporal distribution, physiological activity or individual growth rates (Somero 2005, Thackeray et al. 2016, Scranton & Amarasekare 2017, Madrigal-González et al. 2018), among many other things. Here, the effects of temperature on species’ geographic distributions and seasonal and diel activities are of particular interest since variation in these aspects can have dramatic consequences for their ecological performance and persistence (Edwards & Richardson 2004, Schweiger et al. 2008, Rader et al. 2013). The ongoing climate changes are drastically modifying the spatial and temporal organization of biodiversity (Chapin III & Diaz 2020), which is leading to spatial and seasonal decouples of interacting species (Sheldon et al. 2011) and, thus, to the disruption of food webs and ecosystem services (Román-Palacios & Wiens 2020). Ecologists and climatologists have accumulated a large amount of evidence on these effects during recent decades, which are especially relevant for ectotherms (Paaijmans et al. 2013). Despite this evidence, how temperature responses integrate across different species’ aspects is largely unknown. To obtain this knowledge is crucial because incongruous responses can lead to incompatible adjustments to temperature changes along biological aspects, compromising the species’ performances under climate change.

Delimiting the actual effect of environmental temperature on the distribution and abundance of species may become difficult when other variables that are either spatially or temporally correlated with temperature are considered simultaneously. For instance, the latitudinal distribution of species in the Northern Hemisphere is associated with historical events and dispersal limitations, whose effects generate geographical patterns that can be confounded with those of temperature variations (Araújo et al. 2008, Hortal et al. 2011, Calatayud et al. 2016, 2019). Similarly, the apparent relationships between temperature and either seasonal or diel activities may be indeed conditioned by life-history constraints related with the time required to complete individual development, species’ voltinism, the phase in which overwintering occurs, photoperiod limitations, light requirements, and the reliance on solar radiation independently on the environmental temperature (Bradshaw & Holzapfel 2007, 2010, Teder 2020). Hence, assessing the predictive value of temperature in accounting for the spatial and temporal variations in species occurrence and abundance would require considering any alternative variables that could play a significant role in these variations.

Experimental setups can help unravel the “true” role of temperature in driving geographical, seasonal and diel patterns for some model organisms while controlling for other variables (Angilletta 2009). However, experiments based on artificial thermal gradients can subject individuals to new and unrealistic stress conditions, thereby providing overestimated projections of species responses (Guo et al. 2020). Alternatively, one could explore the contribution of temperature using observational data where the variations in temperature and other complementary predictors are decoupled. For example, the effects of temperature and solar radiation can be teased apart using diel activity from consecutive days that showed substantial variations in temperature (*i.e*., while presenting almost equal sunlight incomings). Similarly, the effects of temperature and day length can be teased apart using seasonal data along steep temperature gradients, with nearly equal day lengths (such as *e.g*., elevational gradients). Finally, the role of temperature in determining the species’ distribution can be assessed by comparing geographical areas with different temperature regimens. That is, if temperature is an important variable, we should find similar responses under different background temperatures.

The relevance of temperature in accounting for the spatiotemporal variation in species occurrence and abundance may thus be estimated from observational data, comparing the results from including or not alternative predictors to account for complementary causal factors. Temperature will stand out as a relevant factor across different biological scales if its association with several species’ responses is high throughout different spatiotemporal dimensions, but also if such responses are congruent across dimensions. The congruence in thermal responses to diel, seasonal and geographical gradients would support the universal and homogeneous role of temperature in delimiting the occurrence and abundance of species. Note here that expectations are that different mechanisms are behind the response to temperature variations associated with geography, seasonality and diel rhythms. For instance, daily temperature variations should also be related to changes in light or other environmental factors that can generate behavioural, endocrine, and physiological diel rhythms (Levy et al. 2019). In contrast, responses to seasonal temperatures should be associated with the annual rhythms and the need to synchronize life history phases with seasonal variations in climate (Saunders 2020). On the other hand, responses to geographical variations in temperature should relate to local adaptation processes acting at the population level, and likely involving the above-mentioned individual tolerances and ontogenetic timing, as well as other essential species attributes (Sunday et al. 2019).

Despite these differences, a certain level of congruence in the responses would indicate the consistent role of temperature as a holistic and predictable driver of key biological aspects. Such congruence would be evident, for example, if species occurring in colder regions are also active during colder periods of the year and at colder hours of the day in areas of milder climate. Such hypothesized thermal congruence is fundamental to respond adequately to global warming, as decoupling responses across different spatiotemporal gradients may expose local populations to critical temperatures, thus compromising their long-term persistence. For instance, if seasonal and diel responses to temperature are decoupled, species might not be able to adjust seasonal cycles as much as it would be necessary to prevent individuals from facing critical temperatures during diel activities. Following this line of evidence, studying the congruence of thermal responses across evolutionary lineages is also important because a marked phylogenetic signal in thermal niches would also point to the relevance of temperature changes. That is, if thermal adaptations are evolutionarily conserved, species might present limited ability to modify their thermal responses, being unable to cope with climate warming. Should this be true, phylogenetic biases in the potential effects of climate warming would be also expectable. Despite the relevance of studying the consistency of thermal responses across spatiotemporal gradients and evolutionary lineages, integrative studies on this topic are lacking.

Here we study the thermal responses associated with geographical, seasonal, and diel temperature variations using several temperate dung beetle species as a model system. Dung beetles are capable to self-regulate their body temperature and produce heat depending on their body size (Verdú & Lobo 2008, Verdú et al. 2012) a physiological adaptation directly linked to the need of a quick dispersal response to exploit an ephemeral resource. In addition, they feed on cattle from domestic and wild animals, participating in nutrient cycling and seed dispersion (Nervo et al. 2017, Milotić et al 2019), and thus, providing important ecosystem functions. These characteristics make dung beetles an ideal and important group to study thermal responses.

Specifically, we evaluated the responses of dung beetles to changes in temperature associated to: (i) diel rhythms across three consecutive days with contrasted temperatures; (ii) seasonal rhythms across six sites located at different elevations; and (iii) geographical ranges along five river basins in the Iberian Peninsula (Fig. 1). We hypothesized that if temperature is the main factor determining the activity and distribution of dung beetles, its effect should be observed along the three considered spatiotemporal gradients, and its relevance would be higher if the effects of other alternative and/or complementary factors are low. Furthermore, congruence in the different species’ thermal responses to diel, seasonal and geographical changes would be expected if the importance of temperature is independent of the spatiotemporal context. On the contrary, a low explanatory capacity of temperature and a lack of congruence in its effects across the three spatiotemporal gradients would support a limited and dissimilar role of temperature depending on the spatiotemporal context. Finally, if species are evolutionarily limited to adapt to new thermal regimens, we expect thermal niches to be phylogenetically conserved.

**Figure 1.**
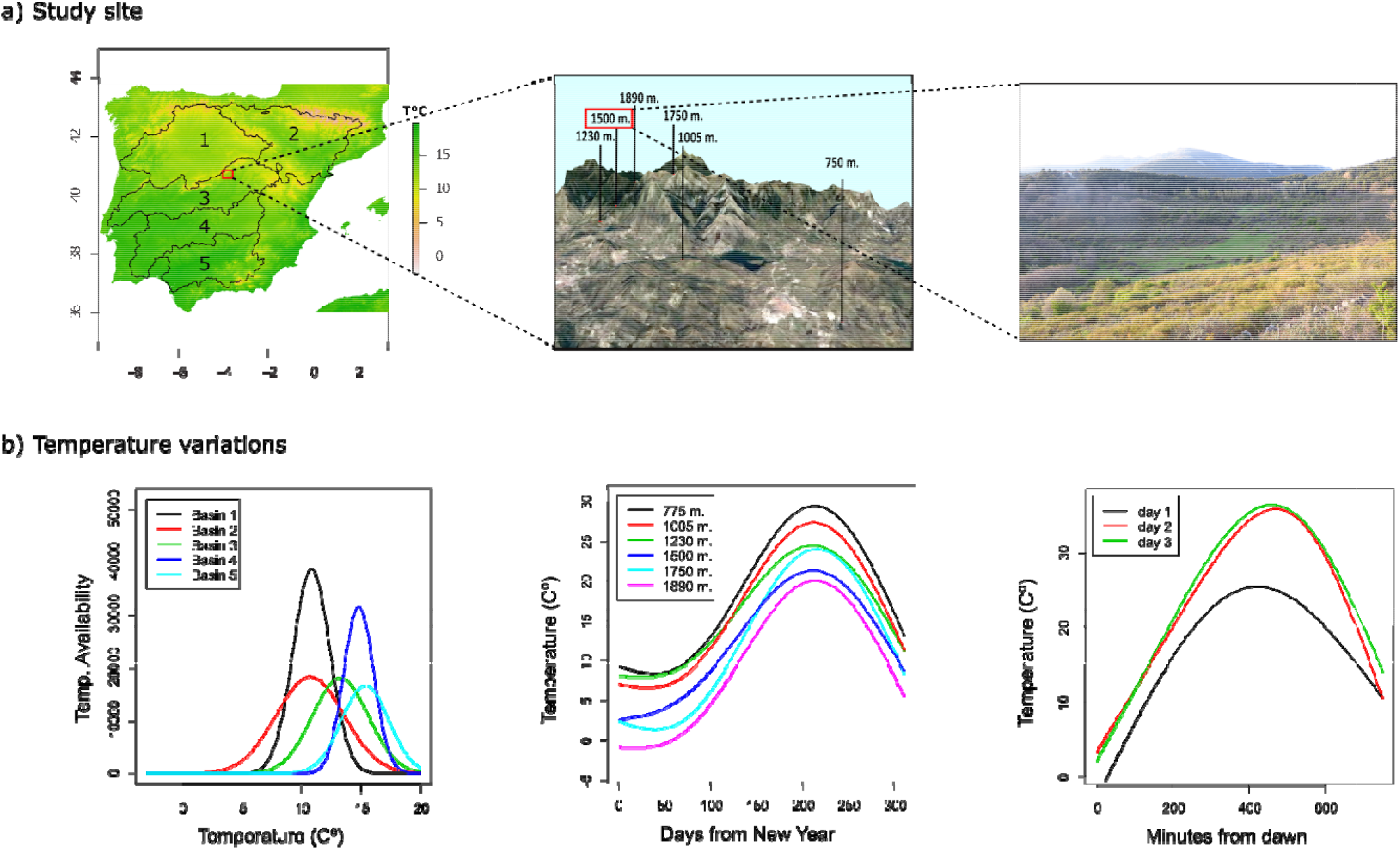
a) The areas of study for the geographical, seasonal and diel datasets (from left to right). Red squares show the position of the following down-scaled study site. b) Temperature variations in study sites. Lines correspond with predictions of general additive models (GAM) of: (i) temperature availability (measured as the number of 10 km^2^ grid cells whose temperature fell within predefined temperature bins) as function of temperature for the geographic dataset (left); (ii) temperature as a function of days from New Year and minutes form dawn for the seasonal and diel datasets respectively. Analyses were computed independently for each basin, for each elevational site and for each day. GAMs explained an average of 0.90 of deviance across all analyses (median = 0.92, ranging from 0.79 to 0.97).

## Material and methods

### Data origin

We use data on 16 Iberian dung beetle species of the family Scarabaeidae (ten from Aphodiinae and six from Scarabaeinae subfamilies). These species were selected because they occurred in at least 10% of the samples of the three datasets considered, covering different spatial and temporal extents (see below). All considered species (Table 1) are of small body size, with body weights far smaller than 1.9 g (0.2 g at most), the threshold from which endothermy is thought to appear in this group of beetles (Verdú et al. 2006). Temperature–occurrence associations for all these species were examined along: (i) five geographical areas of similar extent but different temperature regimes within the Iberian Peninsula (geographical dataset or GD); (ii) six sites placed across a steep elevational range in Central Iberia, and sampled during the same dates but differing in their environmental temperatures (seasonal dataset or SD); and (iii) three consecutive days with similar daily variations but different weather conditions in a single locality near the centre of the same elevational range (diel dataset or DD).

**Table 1.**
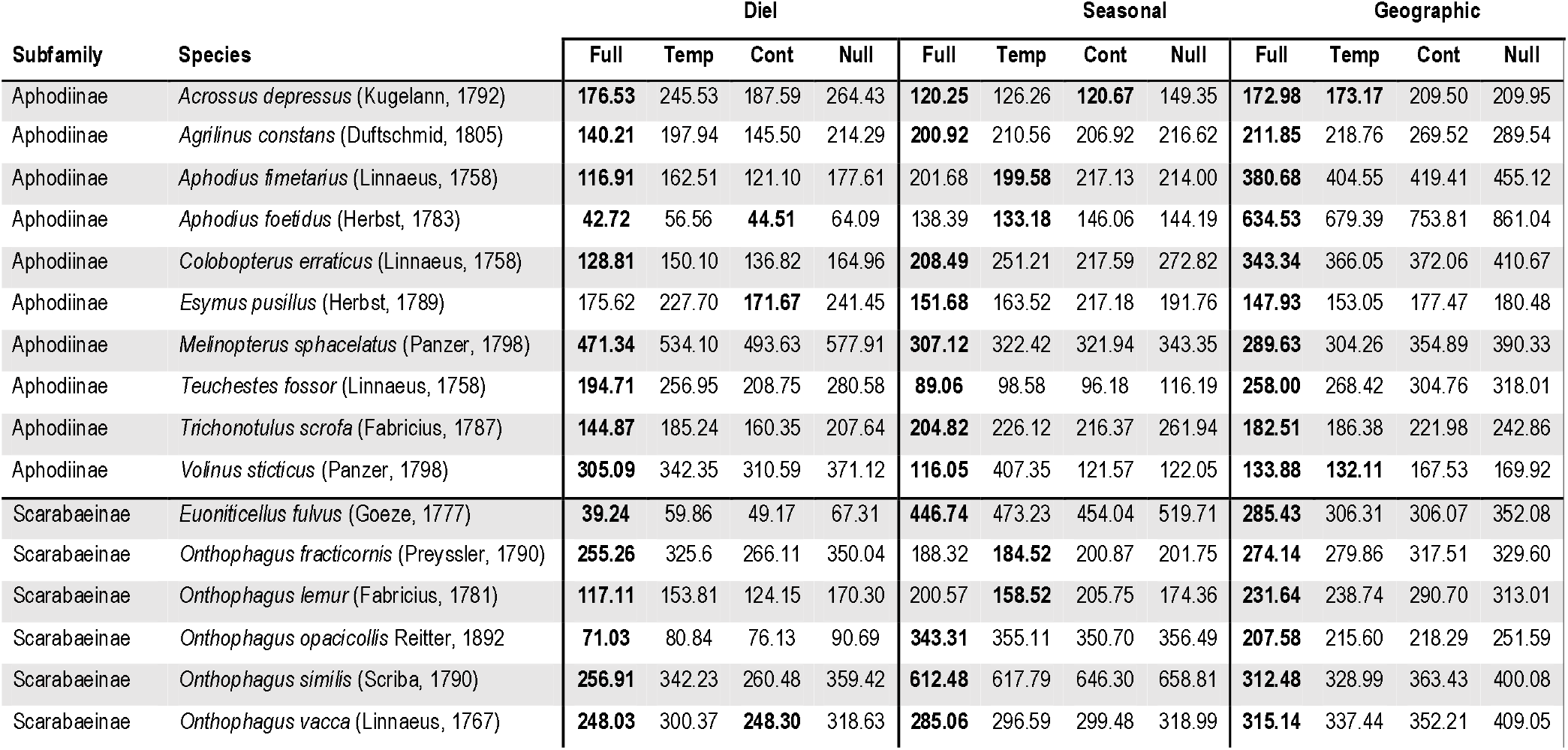
AICc values for the models of each species in each dataset. In all cases, we conducted a complete model (Full) including temperature and the corresponding contrast variables, a model only including temperature (Temp), a model only including contrast variables (Cont), and a null model were no predictor variable was included (Null). Contrast variables were minutes from dawn and its quadratic term for the diel data set; date sine and cosine and their quadratic terms for the seasonal dataset; and temperature availability for the geographic data set. The best models in terms of AICc and the equivalent ones (ΔAICc < 2) are highlighted in bold.

#### Geographical Dataset

The GD is divided in five study areas, corresponding to the major river basins of the Iberian Peninsula (Ebro, Duero, Tajo, Guadiana and Guadalquivir; limits extracted from HydroBASINS data available at www.hydrosheds.org, Lehner & Grill 2013, Fig. 1a). These natural areas were used since their borders correspond with marked geographical accidents, which are expected to act as dispersal barriers. Furthermore, they are relatively similar in extent (areas ranging from 5.6 x 10^4^ to 9.7 x 10^4^ km^2^) and almost follow a latitudinal gradient, hence showing contrasting environmental temperatures (Fig. 1b). In each of these basins, we collected all georeferenced occurrences of the selected species from GBIF (www.gbif.org, accessed May 2020) and additional published sources (Hortal & Lobo 2011). As this kind of data is biased due to historically uneven sampling effort (Lobo et al. 2018), the occurrences were pooled within UTM grid cells of 10 x 10 km spatial resolution. This grain was selected because it corresponds to the effective resolution of most of the occurrence information in the dataset, and it is appropriate to avoid the effects of oversampled localities while retaining a reasonable climatic detail. The frequency of each species’ occurrence data in temperature bins of 1°C (ranging from −3 to 20°C, n=24) was calculated for each river basin (24 x 5 = 120), and these figures were used as dependent variables in the subsequent regression analyses.

#### Seasonal dataset

Six sites along an elevational gradient located in the Sierra de Guadarrama (Central Spain) (Fig. 1a, Espinoza 2016) were used to explore the effect of temperature variations in SD. Elevations range from 755 to 1900 m a.s.l., separating sites approximately 200 m a.s.l.. Each survey site was sampled approximately every three weeks, totalling fourteen times from May 2012 to June 2013. We choose this elevation gradient because these sites show considerable variations in temperature during the whole period of the surveys (Fig. 1b). The sampling protocol in each periodical sample consisted of five pitfall-traps baited with fresh cattle dung and separated around 30 m from each other. Traps were placed in open habitats to avoid potential habitat and shadow effects and were active during 48 h. The individuals of these traps were pooled together, obtaining an estimation of each species’ abundance per elevation site and date (6 x 14 = 84), which were used as response variables in subsequent statistical analyses.

#### Diel Dataset

Temperature effects on diel activity were assessed using dung beetle data from a grassland located next to El Ventorrillo MNCN field station, placed in the Sierra de Guadarrama at an approximate elevation of 1500 m a.s.l. (Fig. 1a). This locality was chosen as it shows a high diversity of dung beetles (between 30 and 40 species belonging to the considered subfamilies; Cuesta & Lobo 2019). We sampled three consecutive days (April 28^th^–30^th^ 2015) that showed contrasting temperatures, with around 8 °C of difference between the mean temperatures of the coldest and the hottest days (Fig. 1b). Each day, ten pitfall traps baited with fresh cattle dung were distributed around a circumference of approximately 50 m. of radius (*i.e*., traps were at least 30 m apart from each other). Since we intended to measure the flight activity during short periods, the bait was introduced into a nylon stocking piece to avoid the stagnancy of beetle individuals within the dung bait along different sampling events. We checked all traps every 30 min. from dawn to dusk (approximately from 7:30 am to 7:00 pm, n=23), collecting all individuals to subsequently identify them in the laboratory. Traps were also checked during the night to discard nocturnal activity. Individuals from the ten traps were pooled together, obtaining an estimation of the abundance of active individuals from each species each 30 min (23 x 3 = 69), which were further used as dependent variables.

### Temperature measures and alternative correlates

Temperature measures were obtained from different standardized methods for each one of the different spatio-temporal scales considered, but trying to maintain a considerable degree of congruence among them. For the *Geographical Dataset*, we obtained mean annual temperatures at a 30 sec resolution from the WorldClim database (see www.worldclim.org, Hijmans et al. 2005). We preferred mean annual temperatures over monthly average figures since the precise seasonal activity over the complete study area was unknown for most of the species. Nevertheless, spring and autumn temperatures (the seasons when phenological peaks occur for most species) were positively correlated with mean annual temperatures (Pearson’s *r* = 0.99 and 0.97, respectively), so we assume that mean annual temperature is a reasonable proxy for both of them.

For the *Seasonal Dataset*, we set up a temperature data logger in each of the elevational points during the whole period of the study. This device was placed in the shadow at one meter from the ground to escape from extreme temperatures due to insolation, mimicking the meteorological stations on which WorldClim data are based on. Temperature was recorded each 10 min. and we used the mean daily temperature when pitfall-traps were active.

In the case of the *Diel Dataset*, temperature measurements were taken using five data loggers placed in the study site just in the centre of the circumference formed by the traps. Data loggers were placed to recover temperature measurements from the different microclimatic conditions available for dung beetles: two at one meter over the ground, in the sun and shadow; another two directly on the ground, also both in the sun and shadow; and one buried at 10 cm depth. Preliminary results showed that the mean temperatures from the data logger placed on the ground in the sun were those that best correlated with the species’ diel activity, so we used these measurements in subsequent analyses. Temperature was recorded each minute, and average temperatures during the 30 min before traps were checked were used as predictors.

As previously stated, the effects of temperature measurements might be overestimated due to its collinearity with other factors with which it shares spatial (in the case of GD) or temporal structure (in the case of SD and DD). We quantified this potential overestimation effect by using different “contrast variables”, alternative predictors which are often partly correlated with temperature but are either measures or proxies of other potential causal factors for dung beetle spatial and temporal responses. These alternative predictors were temperature availability and survey effort in the case of GD, day of the year for SD, and hour of the day in the case of DD. The effect of temperature on the frequency of occurrence (GD) or abundance (SD and DD) that is independent of these contrast variables was assessed as the “pure” effect of temperature variations that is independent of the range of temperatures available (GD), the period of the year (SD), and the hour of the day (DD) (see analytical methods below).

Temperature availability for each basin is the relative frequency of 10 x 10 km UTM cells in each 1°C temperature bin. This variable aims to represent the thermal spectrum available in each basin. Hence, a high explanatory capacity of this variable on the frequency of occurrence of a species would imply that the apparent thermal preference of this species can be simply because its spatial pattern of occurrence mimics the distribution of temperatures in the analyzed basin. Further, the typical correlation between the observed pattern of occurrence of a species and the spatial distribution of survey effort can also generate spurious correlations between species’ frequency and temperature in each basin. This potential source of error was considered here by calculating the relative frequency for each 1°C temperature bin of all dung beetle records included in the formerly mentioned databases and pooled within the 10 x 10 km UTM cells. Nevertheless, we found that this estimation of survey effort and temperature availability were highly and positively correlated in all basins (Pearson’s r ranging from 0.97 to 0.99), since the most frequent temperatures have been also surveyed more often, which implies that the surveys are randomly allocated within the available temperatures. Consequently, we discarded using survey bias as contrast variable, considering that the effect of temperature availability also includes differences in survey effort. In the case of SD data, the day of the year was obtained by first ordering the available dates from the day corresponding to the summer solstice (June 21^th^ = 0 or 360), to subsequently convert these figures into radians and obtaining two circular variables by calculating their cosine and sine values. Thus, the summer-winter oscillation is represented by the cosine of the date and oscillates from 1 to −1, whereas the spring-autumn transition is represented by the sine of the date scale 1 to −1. Finally, the hour of the day (DD data) is simply codified as the number of minutes from dawn.

### Statistical analyses

#### Explanatory capacity of temperature

We first explored the independent capacity of temperature to explain variations in dung beetle data in GD, SD, and DD. For each dataset, we conducted Generalized Linear Regression Models of the relative frequency or the abundance of each species as a function of temperature values. All data coming from the five basins (in GD), the six elevational sites (SD), and the three days (in DD) were considered at the same time in each one of the three models. A curvilinear quadratic function of temperature was included in all the cases to account for the typical unimodal performance curves of ectotherms (Huey & Kingsolver 1989). A negative binomial error distribution for the dependent variable was assumed to avoid overdispersion issues associated with the Poisson error distribution (Blasco-Moreno et al. 2019), and it was related to the set of predictors via a logarithmic link function. It is important to note that we did not include a term in the models to account for the different spatial (*i.e*., basins and elevations) and temporal (*i.e*., days) units. By doing so, we were ignoring other factors that may affect the distribution and activity of dung beetles, besides temperature and the contrast variables analyzed. However, this allows us to tease apart the effects of temperature and the formerly mentioned contrast variables with a spatial and temporal structure while avoiding model overparameterization. Hence, we fitted three supplementary GLMs representing different hypotheses regarding the importance of temperature: (i) a full model where both temperature and contrast variables are included altogether, (ii) a model including only these contrast variables, and (iii) a null model where only the intercept was included. We assumed a linear relationship between the density of occurrence and temperature availability (GD); whereas in SD and DD, we assumed curvilinear relationships between abundance and contrast variables by including a quadratic term of both the number of minutes from dawn, and date sine and cosine. We used a deviance partition approach (Legendre 1993, see also Calatayud et al. 2019 for the same approach) to calculate the deviance explained by each set of variables alone (*i*.*e*., temperature *vs*. contrast variables; herein, total pseudo R^2^) and once accounting for the collinearity with other variables (herein, partial pseudo R^2^). Model performance was assessed using the Akaike Information Criterion corrected for small sample size (AICc).

#### Thermal niche attributes

Deriving thermal niches from occurrence data typically provides a partial description of the whole potential response of the species (Sánchez-Fernández et al. 2012, Saupe et al. 2018). However, occurrence-based thermal niches may nevertheless be characterized by different attributes such as the optimum temperature and niche breadth (Gouveia et al. 2014, Löffler & Pape 2020, Fig. 2). The temperature optimum of each species was assessed by fitting quadratic curves in a GLM and calculating the maxima as their inflection point (see Villén-Pérez & Carrascal 2015 for a similar procedure). Thermal niche breadth was also obtained as the area under the curve of these fitted curves. Fitted values were normalized to reach a maximum value of one to make calculations comparable among datasets and species.

**Figure 2.**
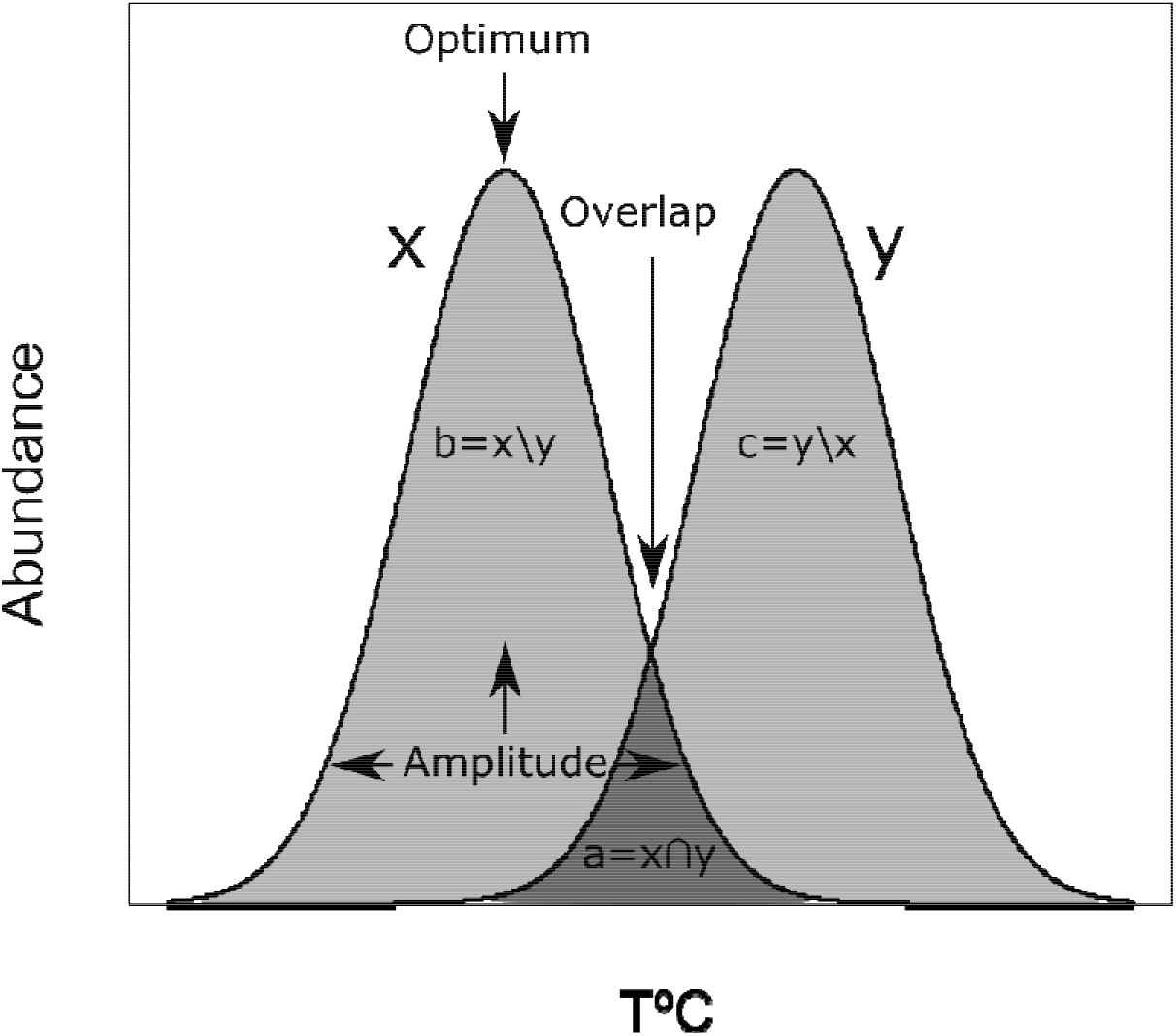
Thermal niche attributes and overlap measure. x and y represent thermal response curves of two species or of a single species in two different study units (*i*.*e*., days, elevation or river basins). From this curve we obtained the optimum temperature and the niche amplitude. Further, we used the overlap between them (a) and the two independent areas (b and c) to calculate the Simpson’s dissimilarity index, as a measure of the congruence between the responses to temperature of the same species at different scales, and of different species within the same scale.

We evaluated the intraspecific dissimilarity in the thermal niches across different spatial and temporal scales, herein called “thermal lability”, using data from the different study units used in each dataset; that is, between river basins, elevation sites, and days (Fig. 2). Thermal lability between pairs of units was measured using the Simpson index as follows:

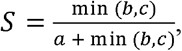

where *a* represents the area under the curves where both curves overlap, and *b* and *c* represent the independent areas under the curves in study units (see Fig. 2). The larger the overlap between the curves obtained at different scales, the smaller the thermal lability will be. We computed this index for all pairs of units in each dataset (*i.e*., for each pair of basins, each pair of elevations, and each pair of days) and then considered the maximum dissimilarity among all pairs from the same dataset, as this measure will provide a more realistic estimate of the potential thermal lability of each species.

#### Congruencies in thermal niches

The congruence in the thermal niches of the different species derived from the three datasets (*i.e*., GD, SD, and DD) was assessed using Spearman rank correlations between the deviance explained by temperature (*i.e*., both for the total and partial pseudo R^2^s), as well as the obtained temperature optima, thermal niche breadths and thermal labilities for each pair of datasets. In addition to these descriptors, we explored the congruence in the overlap of thermal niches estimated from different datasets. To do this, we examined whether interspecific thermal niche dissimilarities were correlated between the different datasets. We computed dissimilarities between the models’ normalized fitted values where the temperature was the only explanatory variable using the Simpson index as previously explained, but in this case between pairs of species (see also Fig. 2). By doing so, we created a thermal niche pairwise dissimilarity matrix for each dataset. Then, we conducted Mantel tests based on Spearman’s ρ correlation coefficient to assess the relationship between dissimilarity matrices obtained from the different datasets. Significance was evaluated by comparing observed ρ coefficients with 999 null values obtained by permuting the dissimilarity matrices.

#### Phylogenetic signal

The potential lability of thermal niches shall be also assessed from an evolutionary point of view. In this sense, a marked phylogenetic signal would indicate both potential evolutionary constrains for temperature variation responses, and phylogenetically-structured effects of global warming. We reconstructed a Bayesian phylogenetic hypothesis for 18 species present in our datasets based on two mitochondrial (COI and COII) and one nuclear markers (28S RNA, see Appendix S1 for details on phylogenetic reconstruction). DNA markers were sequenced for this study and retrieved from Genbank (Table S1, accessed in June 2016). Pagel’s λ test (Pagel, 1999) and Blomberg’s K statistics (Blomberg et al. 2003) were used to explore the phylogenetic signal in the five variables considered (total and partial deviance explained by temperature, temperature optimum, thermal niche breadth, and thermal lability)., Significance for Pagel’s λ was assessed with a likelihood ratio test comparing the negative log likelihood obtained from the original tree topology with the negative log likelihood from a topology transformed to remove the signal (*i.e*., λ = 0). In the case of Blomberg’s K, we tested for significance by randomizing the labels of the phylogenetic tips and comparing observed and random K values. Finally, we also investigated for phylogenetic signal in the thermal niche dissimilarities for each dataset. To do so, Spearman correlations between thermal dissimilarities and phylogenetic distances were used, assessing significance by comparing observed correlations with null values where the labels of the tips of the phylogeny were randomized. In all cases where tip labels were randomized, p-values were calculated as the proportion of null values being equal or higher than observed values.

All analyses were conducted in R environment (R Core Team 2020), using the *AICcmodavg* package (Mazerolle 2019) to calculate AICc values, the function “sintegral” as implemented in the *Bolstad2* packed (Curran 2013) to assess areas under the curves, the *vegan* package (Oksanen et al. 2019) for the Mantel tests, and the *phytools* package (Revell 2012) to calculate Pagel’s λ and Blomberg’s K.

## Results

There is an evident gradient in the explanatory relevance of temperature towards higher relevance at progressively larger scales (*i.e*., geographical > seasonal > diel). Model selection revealed that the full model, including temperature and contrast variables, was the most parsimonious for most species in most datasets (Table 1). As exceptions to this general pattern, in the geographical dataset, the model only including temperature was equivalent to the full model (according to AICc) for one species, and it was also the best supported for another species. In the seasonal dataset, the model only including temperature was the best supported for four species, whereas the model only including contrast variables was equivalent to the full model for just one species. Finally, the model including minute from dawn in DD data was equivalent to the full model for only two species and even better for one species (Table 1). In general, the total deviance explained by the models including temperature and contrast variables was considerably high (mean pseudo-R^2^s = 0.62, 0.63, and 0.77; ranges = 0.51-0.75, 0.38-0.86, and 0.64-0.86, respectively for GD, SD, and DD; see Fig. 3). Partial regressions revealed that the effects of temperature and contrast variables largely overlap, being the deviance independently explained by temperature considerably low (see Fig. 3). Interestingly, the percentage of deviance explained by temperature decreased from the geographical (mean pseudo-R^2^s = 0.33; range 0.13–0.48), to the seasonal (0.19; 0.05–0.36) and diel datasets (0.08; 0.01– 0.20) (see Fig. 3).

**Figure 3.**
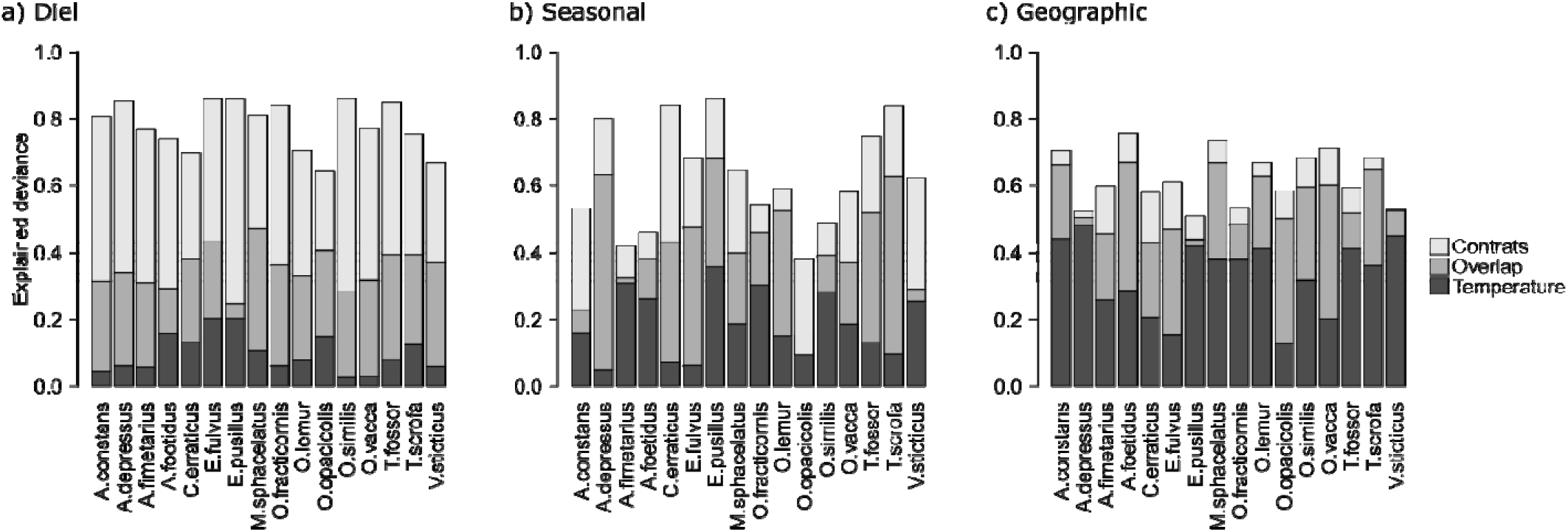
Partial regression results. The deviance explained by temperature alone, the contrast variables alone, and the overlap between them is shown. The contrast variables were minutes from dawn and its quadratic term for the diel data set (a); date sine and cosine and their quadratic terms for the seasonal data set (b), and temperature availability for the geographic data set (c).

Thermal niche attributes derived from the different datasets showed little congruence. Neither the pseudo R^2^ explained by temperature alone nor the total pseudo R^2^ were positively and significantly correlated between any pair of datasets, and none of the thermal niche attributes were significantly correlated between the three considered datasets (Table 2). Moreover, Mantel tests showed that interspecific niche dissimilarities were not correlated among the three studied spatiotemporal scales (Table 2). Finally, we did not find phylogenetic signal for any of these variables in any of the datasets, except in the case of niche breadth for the diel dataset (Table 3).

**Table 2.**
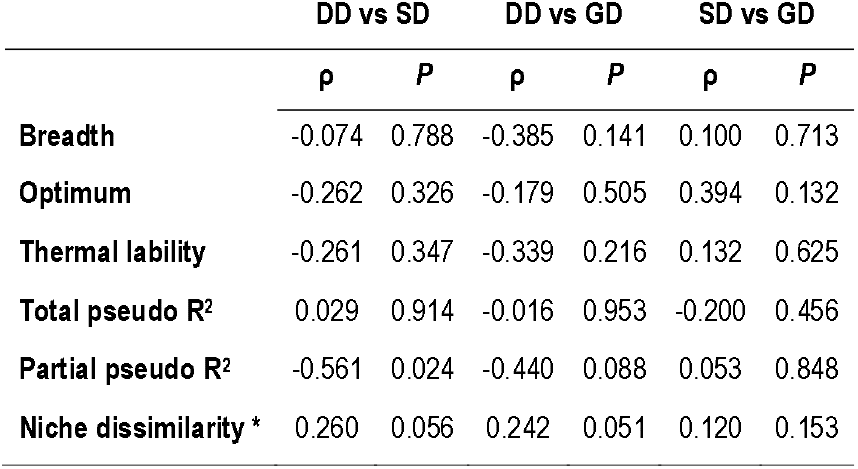
Spearman’s ρ correlation coefficients and P-values between the considered thermal niche attributes measured by the three studied datasets are detailed. DD: Diel dataset. SD: Seasonal dataset. GD: Geographical dataset. * Results based on Mantel test.

|  | DD vs SD |  | DD vs GD |  | SD vs GD |  |
| --- | --- | --- | --- | --- | --- | --- |
| | $\rho$ | *P* | $\rho$ | *P* | $\rho$ | *P* |
| **Breadth** | -0.074 | 0.788 | -0.385 | 0.141 | 0.100 | 0.713 |
| **Optimum** | -0.262 | 0.326 | -0.179 | 0.505 | 0.394 | 0.132 |
| **Thermal lability** | -0.261 | 0.347 | -0.339 | 0.216 | 0.132 | 0.625 |
| **Total pseudo R<sup>2</sup>** | 0.029 | 0.914 | -0.016 | 0.953 | -0.200 | 0.456 |
| **Partial pseudo R<sup>2</sup>** | -0.561 | 0.024 | -0.440 | 0.088 | 0.053 | 0.848 |
| **Niche dissimilarity \*** | 0.260 | 0.056 | 0.242 | 0.051 | 0.120 | 0.153 |

**Table 3.**
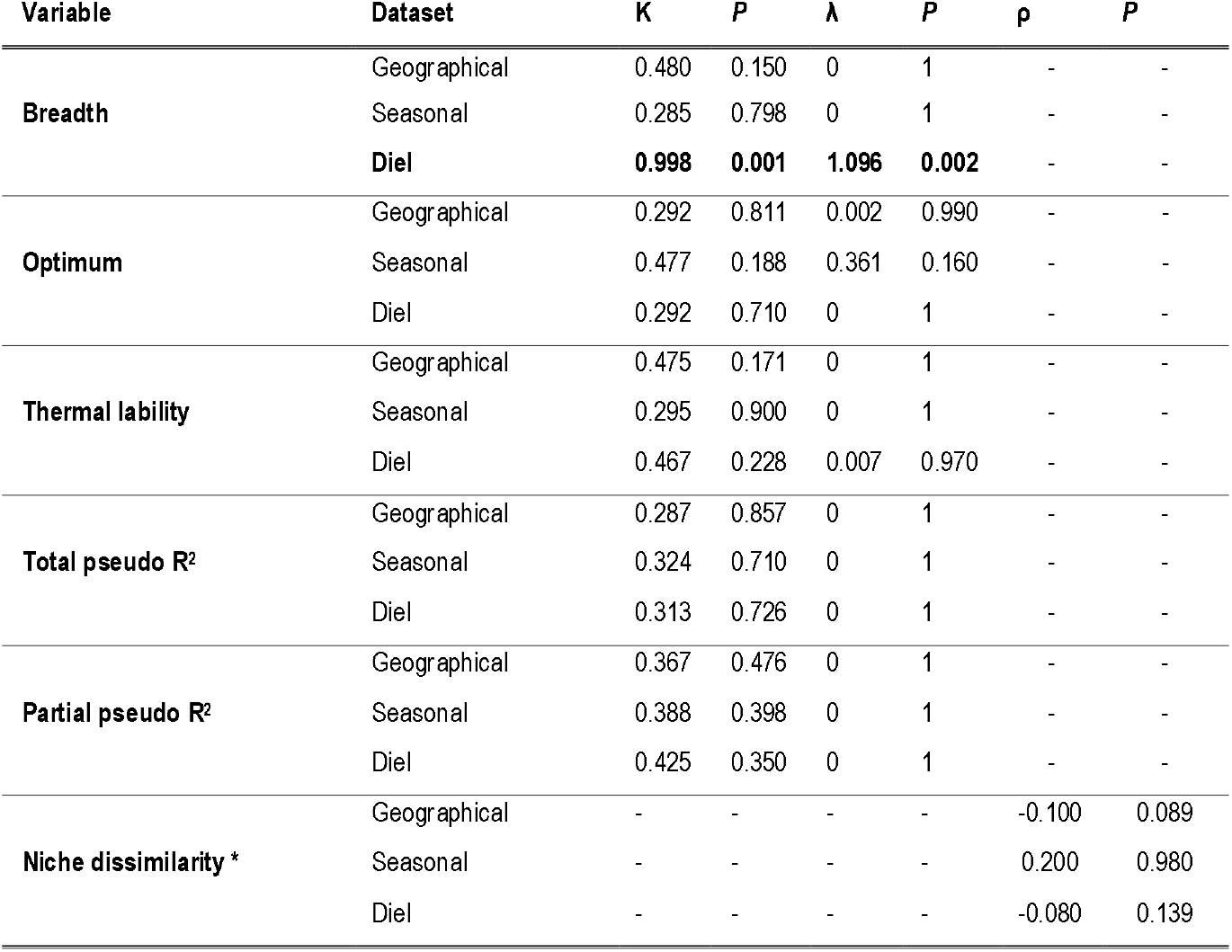
Phylogenetic signal in thermal niches attributes for the three studied datasets (*i.e*., geographical, seasonal and diel). Significant variables are highlighted in bold. * Results based on Mantel test.

| Variable | Dataset | K | P | $\lambda$ | P | $\rho$ | P |
| --- | --- | --- | --- | --- | --- | --- | --- |
| **Breadth** | Geographical | 0.480 | 0.150 | 0 | 1 | - | - |
|  | Seasonal | 0.285 | 0.798 | 0 | 1 | - | - |
|  | **Diel** | **0.998** | **0.001** | **1.096** | **0.002** | - | - |
| **Optimum** | Geographical | 0.292 | 0.811 | 0.002 | 0.990 | - | - |
|  | Seasonal | 0.477 | 0.188 | 0.361 | 0.160 | - | - |
|  | Diel | 0.292 | 0.710 | 0 | 1 | - | - |
| **Thermal lability** | Geographical | 0.475 | 0.171 | 0 | 1 | - | - |
|  | Seasonal | 0.295 | 0.900 | 0 | 1 | - | - |
|  | Diel | 0.467 | 0.228 | 0.007 | 0.970 | - | - |
| **Total pseudo R<sup>2</sup>** | Geographical | 0.287 | 0.857 | 0 | 1 | - | - |
|  | Seasonal | 0.324 | 0.710 | 0 | 1 | - | - |
|  | Diel | 0.313 | 0.726 | 0 | 1 | - | - |
| **Partial pseudo R<sup>2</sup>** | Geographical | 0.367 | 0.476 | 0 | 1 | - | - |
|  | Seasonal | 0.388 | 0.398 | 0 | 1 | - | - |
|  | Diel | 0.425 | 0.350 | 0 | 1 | - | - |
| **Niche dissimilarity \*** | Geographical | - | - | - | - | -0.100 | 0.089 |
|  | Seasonal | - | - | - | - | 0.200 | 0.980 |
|  | Diel | - | - | - | - | -0.080 | 0.139 |

## Discussion

Our results show that the spatial and temporal responses of the studied species show large associations to contrast variables besides temperature, but also that temperature controls to dung beetle occurrence may increase towards larger temporal and spatial scales. This contrasts with our preliminary expectations of a high importance of temperature for dung beetle occurrence and activity based on the known basal ectothermic physiology of the considered species. Further, thermal niches were incongruent across scales for the studied species and lacked phylogenetic signal, indicating that thermal adaptations are highly variable both between and within species.

The generally low partial effects of temperature found in our study lead to two important conclusions: (i) the abundance, distribution, and activity of dung beetles are controlled by other factors different from temperature, which are at least partially represented by the *ad hoc* contrast variables used here; and (ii) dung beetle species must have biological mechanisms that provide them with the plasticity required to cope with the temperature variations associated to each spatiotemporal context. Thermoregulation and body heat gain are intimately linked to solar radiation in ectotherms (Angilletta 2009). Indeed, empirical evidence suggest that solar radiation is associated with dung beetles’ body temperatures (Bartholomew & Heinrich 1978) and temporal variations in their abundance and species richness (Lobo et al. 1998). Hence, it is likely that this factor is a key environmental control of the diel activity of dung beetles. Regarding annual rhythms, photoperiod seems to be a crucial environmental cue regulating insects’ seasonality (Nijhout 1994, Bradshaw & Holzapfel 2007). This is likely the case for dung beetles, given the relatively weak effects of temperature on their phenology found in our study. Also, the different life-history phases of an insect need to be synchronized seasonally, and these require a minimum amount of time to complete. The development of a dung beetle individual requires from 30 to 80 days depending on the species (Christensen & Dobson 1977, Romero-Samper & Martín-Piera 1995, 2007, Arellano et al. 2017), a time that determines key life-history characteristics such as the number of generations per year or the overwintering phase. These developmental constraints are therefore hard to modify without major evolutionary changes (Teder 2020), thereby limiting the effects of environmental temperature on the seasonal abundance and occurrence of dung beetle species. Finally, many factors contribute to shaping the geographical distribution of dung beetle species, including dispersal limitations (Lobo et al. 2006), historical events (Hortal et al. 2011), or the response to other environmental variables such as precipitation, soil, habitat, or trophic preferences (Hanski et al. 1991, Hortal et al. 2001, Lobo & Martín-Piera 2002). It is important to note that we have not quantified the effects of these variables explicitly, so their inclusion could further weaken the pure effect of temperature.

Regardless of the effects of alternative factors, it seems accurate that dung beetles have mechanisms to withstand marked temperature variations, especially those associated with diel and seasonal rhythms. Given the nature of our data and analyses, these mechanisms can be operating either at the population level, at the individual level, or both. At the population level, a high genetic diversity linked to large phenotypic variability can produce the apparently labile thermal responses. That is, as individuals are sorted in time and/or space according to their environmental adaptations, population(s) formed by individuals with different thermal preferences would show a certain level of thermal independency. This mechanism seems more plausible to explain results in the geographical datasets, where river basins can act as dispersal barriers, limiting gene flow and enhancing local adaptation to different temperature regimens (Lenormand 2002). However, it seems less likely that this phenotypic variability alone is responsible for the responses to diel and seasonal temperature variations, where a high gene flow is expected between the individuals and populations that are active at different elevations or days. Physiologically plastic responses allowing individuals to be active at different temperatures seem a more plausible mechanism in this case (Crispo 2008). In any case, these two potential mechanisms (phenotypic variability and individual plasticity) are in agreement with the observed lack of phylogenetic signal on species responses to temperature across scales, which indeed suggests a lack of thermal niche conservatism (Gilbert & Miles 2019). The relative contribution of population phenotypic variability and individual plasticity remains elusive, calling for further studies directed to unravel the detailed mechanisms behind the diverse responses to temperature found in our study.

Be that as it may, the effects of temperature were significant and not negligible, being larger for species distribution than for seasonal activity, and even smaller for diel activity. The increasing importance towards larger scales may be related to the fact that the effects of temperature on the studied biological aspects are nested. That is, the occurrence in a given location would entail that a species holds the adaptations required to maintain a stable population there, which include physiological and/or behavioural adaptations to cope with the seasonal temperature variations that occur in that locality. In the same way, a population with adults active during a given period of the year should present adaptations to handle the daily temperature variations happening during the days when adults are active. Hence, the hierarchically cumulative effects of temperature across these biological scales may explain why temperature becomes more important for geographic distributions than for temporal activities. Ascertaining the plausibility of this idea requires further investigation of intraspecific responses to daily temperature variations across seasons and seasonal temperature responses throughout different populations placed across the species’ geographic distribution.

Perhaps the most interesting of our results is the lack of congruence in the realized thermal niches across the studied species and spatiotemporal contexts. This means that, for instance, species occurring in colder regions do not appear in colder months nor at colder hours of the day in other regions. This somehow counterintuitive result could be related to the uneven relevance of the alternative variables for the different species and spatiotemporal contexts, which facilitates the decoupling of the thermal responses associated with the distribution and activity of dung beetles. It is likely that the processes involved in adult movements, life-history cycles, and population maintenance are differently regulated by temperature, despite of their nested nature. In other words, our results suggest that species have multidimensional thermal niches, where each critical biological aspect responds to temperature along a different dimension. Therefore, rather than exerting a universal effect, temperature plays multiple roles in a species’ biology and metapopulation dynamics. This lack of congruence, together with the low independent effects of temperature found in our deviance partition analyses, indicates that estimates of thermal niches will be, in general, inaccurate and context-dependent. This calls from being particularly cautious when using responses measured at different scales as proxies for future responses to climate change. Overall, our results show the difficulties in estimating general thermal niches of species, challenging forecasts of species future dynamics under climate warming based on unidimensional thermal niches (Gvoždik, 2018).

The partial control of temperature on the activity and distribution of dung beetles may be both a blessing and a curse regarding the effects of climate warming. On the one hand, the apparent thermal lability suggests that temperature increases should not strongly modify neither diel and seasonal activities nor the geographic distribution of dung beetles, likely preventing mismatches with interacting species and the subsequent food chain perturbations. This assumption would contradict the results of studies suggesting moderate or even large effects of climate change on dung beetle distributions (Dortel et al. 2013, Menéndez et al. 2013, Holley & Andrew 2019). On the other hand, the diel, seasonal, or geographical adjustments are among the fastest responses to climate warming (Levy et al. 2019, Duchenne et al. 2020). However, our results suggest that the response towards temperature variations is relatively independent at each spatiotemporal scale. This entails that adjustments to temperature requirements may not be coordinated across key biological aspects. Hence, adjustments to fulfil the temperature requirements for one biological aspect may result in detrimental effects on other aspects, thereby reducing individual and population performance as, *e.g*., seasonal adjustments may expose individuals to inadequate temperatures during diel activity. In the worst-case scenario, the incapacity of species to adjust their temperature requirements by modifying diel, seasonal, and geographical patterns at convenience will increase the likelihood of local extinctions when the individuals are exposed to critical temperatures in their daily or yearly periods of activity. Paradoxically, the partially weak effects of temperature we found may have serious consequences for climate warming if temperature regulates important aspects of species’ biology in divergent ways (Tsai et al. 2020).

Overall, our results show that temperature may be less important than other factors in determining dung beetle activity and distribution. Further, the incongruences in thermal niches estimated from the geographic distribution and seasonal and diel activities show the complex effects of temperature on key species aspects, pointing to a truly multidimensional nature of thermal niches. Together with the partially weak control of temperature on species activity and distribution, these incongruences may difficult fast responses to climate warming, potentially exposing individuals to critical, or at least inadequate, temperatures and reducing individual and population’s fitness.

## Supporting information

### Appendix S1

Genomic DNA was extracted from each individual using the BIOSPRINT 15 DNA Kit (Qiagen), following standard manufacturer’s protocols for blood, and resuspended in 100 μl of buffer AE. We used COI Sca F, COI Sca R, COII am Sca and COII B 605 Sea (Villalba et al. 2002) and the universal 28S a y 28S 5b primers to amplify fragments of the mitochondrial cytochrome oxidase I (COI), the cytochrome oxidase II (COII) and the 28S genes. Amplifications for all gene fragments were performed in a 50 μl reaction containing 39.7 μl of H_2_O, 5 μl of 10x PCR buffer, 1 μl of dNTP mix (10 mM), 0.5 μl of each primer (10 μM), 0.3 μl of AmpliTaq^®^ DNA polymerase (Applied Biosystems) and 3 μl of DNA template. Thermocycling conditions consisted of an initial denaturing step at 94 °C for 4 min, followed by two cycles: (i) a precycle of 5 amplification cycles of 94 °C for 45 sec, 40 °C for 1 min and 72 °C for 1 min, and (ii) a cycle of 35 amplification cycles of 94 °C for 45 sec, 44 °C for 1 min and 72 °C for 1 min, followed by a final elongation step at 72 °C for 10 min and a rapid thermal ramp down to 4 °C. For all reactions, the presence of amplicons of the expected sizes was checked by electrophoresis on a 0.8 % agarose gel. PCR products were purified with the ethanol-precipitation method (Sambrook et al., 1989). Sequencing was performed by Secugen S.L. (Madrid, Spain), using BigDye^®^ and the automated ABI PRISM 3730xl DNA Analyzer. Sequence chromatograms were read and contigs assembled using Sequencher version 4.7 (Gene Codes Corporation, Ann Arbor, MI). All new sequences were deposited in GenBank (see accession numbers in Table S1).

Sequences were aligned in CLUSTALW and MUSCLE, followed by visual inspection using BioEdit (Hall, 1999). Prior to phylogenetic analysis, jModeltest 2.1.1 (Darriba et al., 2012) was used to choose the best-fit model of nucleotide substitution for each of the four genes, and for combined matrices under the corrected Akaike information criterion (AICc). For the COI and COII, HKY was obtained, while Jukes Cantor for 28S. Phylogenetic analyses were performed in a Bayesian framework using BEAST v 2.4 (Drummond and Rambaut, 2007). We established 3 calibrations points based on Ahrens et. al (2014), setting uniform priors with lower and upper boundaries. The calibrations represent the basal split of the following taxa: Aphodiinae (58.7 – 55.8 Million years ago), *Aphodius* (37.2 – 33.9 Mya) and Scarabaeinae (92 – 83.5 Mya). For the age of the rest of the nodes, we set a LogNormal relaxed molecular clock for each gene and let the software estimate the rate from the priors. The MCMC chain ran for 100.000.000 steps, sampled every 10.000 steps. Posterior distribution of all the parameters were checked using Tracer, as well as all ESS values being above 200. We built the tree using Tree Annotator, using the Maximum Clade Credibility implemented method after discarding the first 25% samples as a burn-in.

---

**Fig. S1.**
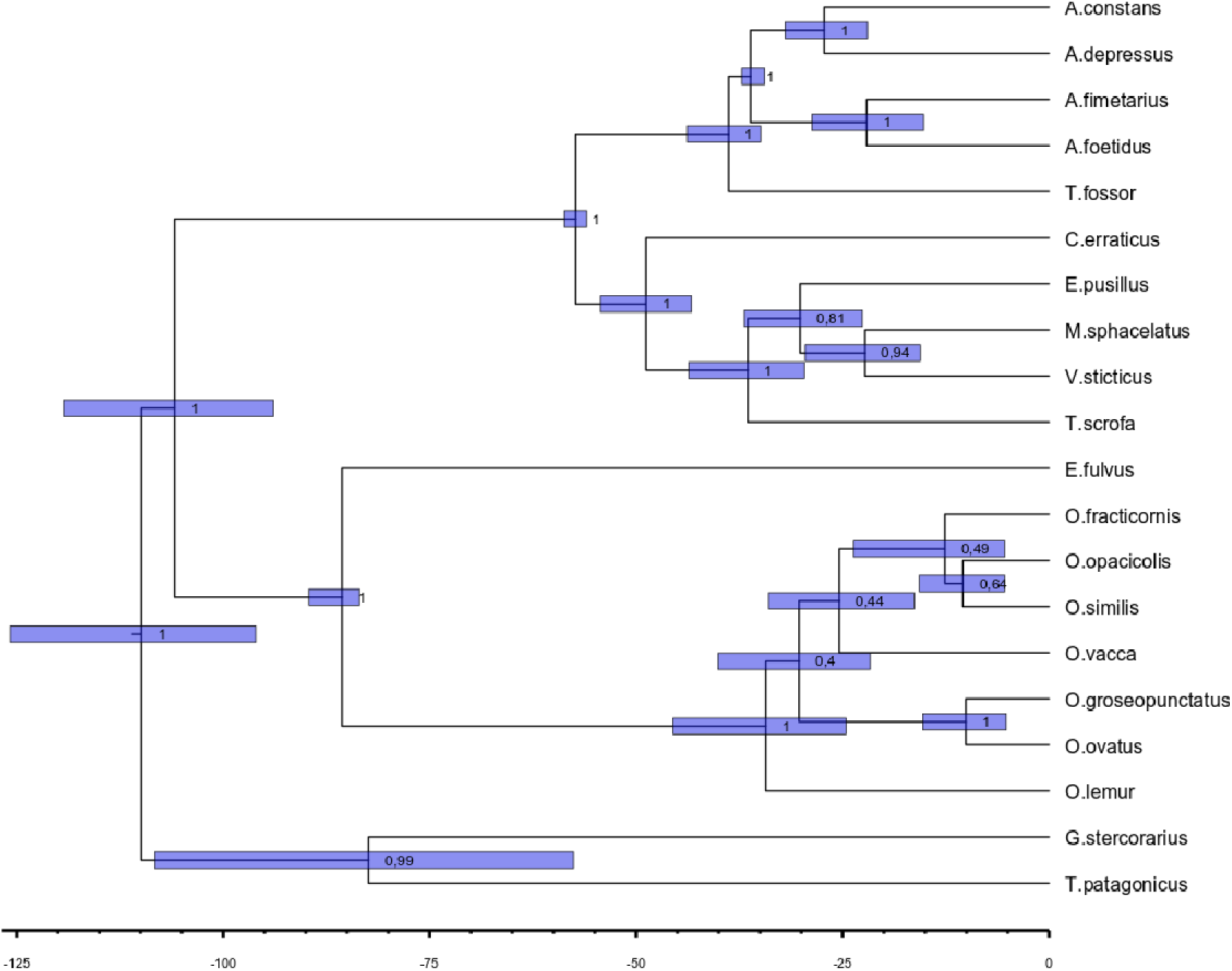
Bayesian phylogenetic hypothesis for the studied species. Posterior probabilities are provided. Blue bars represent the 95% credible interval around node ages.

**Table S1.** GenBank accession numbers of the used sequences. Outgroup species are indicated.

| **Species** | **28** | **COI** | **COII** |
| --- | --- | --- | --- |
| *Aphodius constans* | - | AY039372 | AY039372 |
| *Aphodius depressus* | ABXXXX | ABXXXX | ABXXXX |
| *Aphodius erraticus* | ABXXXX | ABXXXX | ABXXXX |
| *Aphodius fimetarius* | ABXXXX | ABXXXX | ABXXXX |
| *Aphodius foetidus* | - | ABXXXX | ABXXXX |
| *Aphodius fossor* | ABXXXX | ABXXXX | ABXXXX |
| *Aphodius pusillus* | ABXXXX | ABXXXX | ABXXXX |
| *Aphodius scrofa* | ABXXXX | ABXXXX | ABXXXX |
| *Aphodius sphacelatus* | ABXXXX | ABXXXX | ABXXXX |
| *Aphodius sticticus* | - | ABXXXX | - |
| *Euoniticellus fulvus* | ABXXXX | ABXXXX | ABXXXX |
| *Geotrupes stercorarius (OUT)* | KP419463 | AY039377 | AY039377 |
| *Onthophagus fracticornis* | ABXXXX | - | - |
| *Onthophagus grossepunctatus* | ABXXXX | AY039347 | AY039347 |
| *Onthophagus lemur* | ABXXXX | AY039353 | AY039353 |
| *Onthophagus opacicollis* | - | ABXXXX | ABXXXX |
| *Onthophagus ovatus* | ABXXXX | AY039351 | AY039351 |
| *Onthophagus similis* | ABXXXX | ABXXXX | ABXXXX |
| *Onthophagus vacca* | ABXXXX | AY039359 | AY039359 |
| *Taurocerastes patagonicus (OUT)* | KP419662 | GU984611 | GU984611 |

